# Determining the effects of temperature on the evolution of bacterial tRNA pools

**DOI:** 10.1101/2023.09.26.559538

**Authors:** Vatsal Jain, Alexander L. Cope

## Abstract

The genetic code consists of 61 codon coding for 20 amino acids. These codons are recognized by transfer RNAs (tRNA) that bind to specific codons during protein synthesis. Most organisms utilize less than all 61 possible anticodons due to base pair wobble: the ability to have a mismatch with a codon at its third nucleotide. Previous studies observed a correlation between the tRNA pool of bacteria and the temperature of their respective environments. However, it is unclear if these patterns represent biological adaptations to maintain the efficiency and accuracy of protein synthesis in different environments. A mechanistic mathematical model of mRNA translation is used to quantify the expected elongation rates and error rate for each codon based on an organism’s tRNA pool. A comparative analysis across a range of bacteria that accounts for covariance due to shared ancestry is performed to quantify the impact of environmental temperature on the evolution of the tRNA pool. We find that thermophiles generally have more anticodons represented in their tRNA pool than mesophiles or psychrophiles. Based on our model, this increased diversity is expected to lead to increased missense errors. The implications of this for protein evolution in thermophiles are discussed.

**Significance:** Protein synthesis is a vital biological process; however, our understanding of the impact of environmental factors, such as temperature, on the evolution of the molecular mechanisms involved in protein synthesis is limited. In this study, we investigated the impact of environmental temperature on the evolution of the tRNA pool. Our analyses revealed that heat-loving bacteria (thermophiles) generally have more anticodons represented in their tRNA pool. Based on a simple model of ribosome elongation, this observed increase in tRNA diversity in thermophiles is expected to also increase the frequency of translation errors. We speculate that the increased diversity of the tRNA pool could be due to the decreased efficiency of wobble base pairing at higher temperatures, necessitating more tRNA with exact codon-anticodon pairings. Our findings provide key insights into the role of the environment in shaping the tRNA pool.

## Introduction

Life exists in a wide variety of environments, even those that would be considered extreme relative to the environments humans occupy, ranging from deep-sea hydrothermal vents to arctic regions (Merino et al. 2019). Organisms found in these extreme environments play an important role in the cycling of nutrients and energy, and they are also important for the decomposition of organic matter (Merino et al. 2019). Thus, there is interest in studying the mechanisms and evolutionary adaptations that allow these organisms – broadly referred to as “extremophiles” – to not only survive but thrive in these extreme environments. Of particular interest is the ability of bacteria to thrive in extreme temperature environments.

A key process hypothesized to undergo adaptation to extreme environments is the process of protein synthesis. A large portion of a cell’s resources is dedicated to protein synthesis, with about 50% in growing bacteria cells (Buttgereit & Brand 1995; Russell & Cook 1995). Although the basic process of protein synthesis is conserved across all domains of life, extremophilic organisms are hypothesized to have undergone many adaptations to maintain functional protein synthesis under conditions that would stress their mesophilic counterparts. For example, thermophilic (heat-loving) bacteria are hypothesized to have higher GC% in their protein-coding genes, rRNA, and tRNA as an adaptation to higher temperatures, although the relationship between temperature and GC% of protein-coding genes appears to be due to shared ancestry rather than true adaptation. Relative to mesophiles, thermophiles, and psychrophiles are hypothesized to invest less or more, respectively, in translation resources. Thermophiles are observed to have relatively few tRNA and ribosomal RNA (rRNA) genes encoded in their genomes, while psychrophiles are observed to have a greater number of these genes. Relatedly, previous work found that the degree of codon usage bias varies across psychrophiles, mesophiles, and thermophiles, with psychrophiles having the most biased genomes and thermophiles having the least biased. This is hypothesized to reflect stronger (weaker) selection on translation efficiency at lower (higher) temperatures due to slower (faster) diffusion and chemical reaction rates (Arella et al. 2021; Vieira-Silva & Rocha 2010).

A crucial aspect of protein synthesis is the tRNA pool, making it a likely target for potential adaptations related to environmental factors. Notably, there have been numerous observations made about the apparent impact of environmental temperatures on the evolution of the tRNA pool. Aside from the apparent relationship of GC% and the number of tRNA genes with temperature, adaptation to extreme environments is also thought to occur via tRNA nucleotide modifications. For example, the modified nucleotide pseudouridine Ψ is more prevalent in psychrophilic bacteria, a supposed adaptation to maintain the flexibility of tRNA at colder temperatures. Additionally, the number of anticodons represented is hypothesized to be a possible adaptation related to temperature. Species generally use fewer than all possible 61 anticodons due to wobble matching at the 3rd nucleotide of the codon. Previous work concluded thermophiles have the most diverse tRNA pools (i.e. the most anticodons represented) despite having the fewest number of tRNA genes, on average (Satapathy et al. 2010; Vieira-Silva & Rocha 2010). In contrast, psychrophiles had the least diverse tRNA pools, on average. This observation could have implications for the effectiveness of wobble base pairing as a function of temperature; however, this analysis was based on a limited set of species and failed to account for the shared ancestry of the species.

The structure of the genetic code and the currently established wobble rules are hypothesized to reduce the frequency and the impact of missense errors (Koonin & Novozhilov 2009). Despite this, mRNA translation is far more error-prone than either DNA replication or transcription, with an incorrect amino acid being incorporated into a peptide chain 1 out of every 1,000 to 10,000 codons (Drummond & Wilke 2009; Ogle & Ramakrishnan 2005). A key factor shaping missense error rates are the ratios of cognate to near-cognate tRNA for each codon (Shah & Gilchrist 2010; Mordret et al. 8 2019). Presumably, a more diverse tRNA pool would lead to higher missense error rates, potentially leading to other evolutionary adaptations to either reduce the frequency of missense errors or reduce the impact of missense errors when they occur.

Here, we examine the evolution of the tRNA pools across a phylogenetically diverse set of bacteria. We find that thermophilic bacteria generally have a greater number of tRNA anticodons represented in their genomes compared to mesophiles or psychrophiles. This increase in tRNA diversity is expected to increase the average missense error rate under standard wobble rules. We speculate the increased tRNA diversity may result from a lowered efficiency of wobble base pairing at higher temperatures, leading to fitness benefits for expanding the number of tRNAs represented in the tRNA pool.

## Materials and Methods

### Obtaining tRNA gene copy numbers

We obtained tRNA gene copy numbers (tGCNs) for 1,093 bacteria from a previous study (Diwan & Agashe 2018). The total number of tRNA genes for each bacterium was calculated by summing the tGCNs across all represented anticodons. The tRNA diversity value was calculated by summing the number of unique anticodons represented in the tRNA pool (e.g. a species with tRNA representing all 61 anticodons would have a tRNA diversity of 61). Organisms are categorized as psychrophilic, mesophilic, or thermophilic depending on their optimal growth temperature: psychrophiles prefer below 15°C, thermophiles above 45°C, and mesophiles between 15°C and 45°C (Taylor & Vaisman 2010). Optimal growth temperatures and GC content were taken from a database of optimal growth temperatures (Helena-Bueno et al. 2023).

### Model of translation errors

To estimate codon-specific elongation rates and missense error rates (i.e. rate of incorrect amino acid incorporation per codon), we implemented a mechanistic model that takes in the tRNA gene copy numbers (tGCN) of a species (Shah & Gilchrist 2010). Standard wobble rules were applied for bacteria as previously done. For each codon in a species, a list of cognate, pseudo-cognate, and near-cognate tRNA were identified. A cognate tRNA is defined as a tRNA with an anticodon that either (1) perfectly matches a codon or (2) matches a codon via wobble rules, with either case resulting in the correct amino acid being inserted into the growing peptide chain. Pseudo-cognates are anticodons that code for the same amino acid as the focal codon, but do not follow standard wobble rules. Near-cognates are anticodons that have a single nucleotide mismatch, but do not code for the correct amino acid. For each codon *i*, we used a species tGCN and model of wobble pairing to estimate the cognate elongation rate *R*_*C*_(*i*), near-cognate elongation rate *R*_N_ (i), and the missense error rate 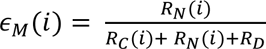, where *R*_D_ is the expected ribosome drop-off rate (i.e. the rate at which premature translation termination occurs) set to be *3*.*146* × *10*^−*3*^*S*^−^*^1^*for all codons. Elongation rates for each species were scaled such that the harmonic mean across all codons was 12.5 amino acids per second as done previously. Further model details can be found elsewhere (Shah & Gilchrist 2010).

### Comparing data across species

Phylogenetic linear regressions were used to test for differences in the tRNA pool as a function of genome-wide GC content and the classification of the species (thermophile, mesophile, psychrophile) while accounting for the shared ancestry of the bacteria (Felsenstein 1985). The phylogenetic tree was taken from a prior study of tRNA pool evolution across bacteria (Diwan & Agashe 2018). This phylogenetic tree was time-calibrated using the penalized likelihood approach implemented in **treePL** with smoothing parameter 0.001 (Smith & O’Meara 10 2012). The final smoothing parameter was determined following a pipeline for **treePL** outlined previously (Maurin 2020). To fit the phylogenetic regressions, linear models comparing data across species were fitted using the function phylolm from the R package **phylolm** with the covariance between species defined by either a Brownian Motion (BM) or Ornstein-Uhlenbeck (OU) model of trait evolution. For comparing across temperature classes, a phylogenetic regression was implemented treating the bacteria’s classification as a discrete variable, with mesophiles serving as a reference class (the intercept term). The best model for each comparison was determined by selecting the model with the minimum Akaike Information Criterion (AIC). In cases where the differences between AIC was < 2 units, indicating relatively small differences in the models, we selected the model with the fewest parameters.

## Results and Discussion

### Evolution of tRNA pool across all bacteria

Our final dataset contained 334 mesophiles, 5 psychrophiles, and 67 thermophiles (Figure 1A). Genome-wide GC% ranged from 26% to 74% with a median of 51%. The diversity of the tRNA pool ranged from 25 to 45 with a median of 41, and the size of the tRNA pool (i.e. the total number of tRNA genes) ranged from 28 to 142 with a median of 48 (Figure 1B,C). We note that the tRNA diversity for mesophiles appears to be bimodal with a small peak at 33 and a larger peak at 43 (Figure 1B, purple). Interestingly, many species with 33 anticodons represented in the tRNA pool are common psychotrophs: bacteria that are able to survive at temperatures < 10 °C, but grow best in the range of mesophilic temperatures. This includes bacteria such as *Psychrobacter arcticus*, *Pseudoalteromonas atlantica*, and *Shewanella halifaxensis*.

**Figure 1.**
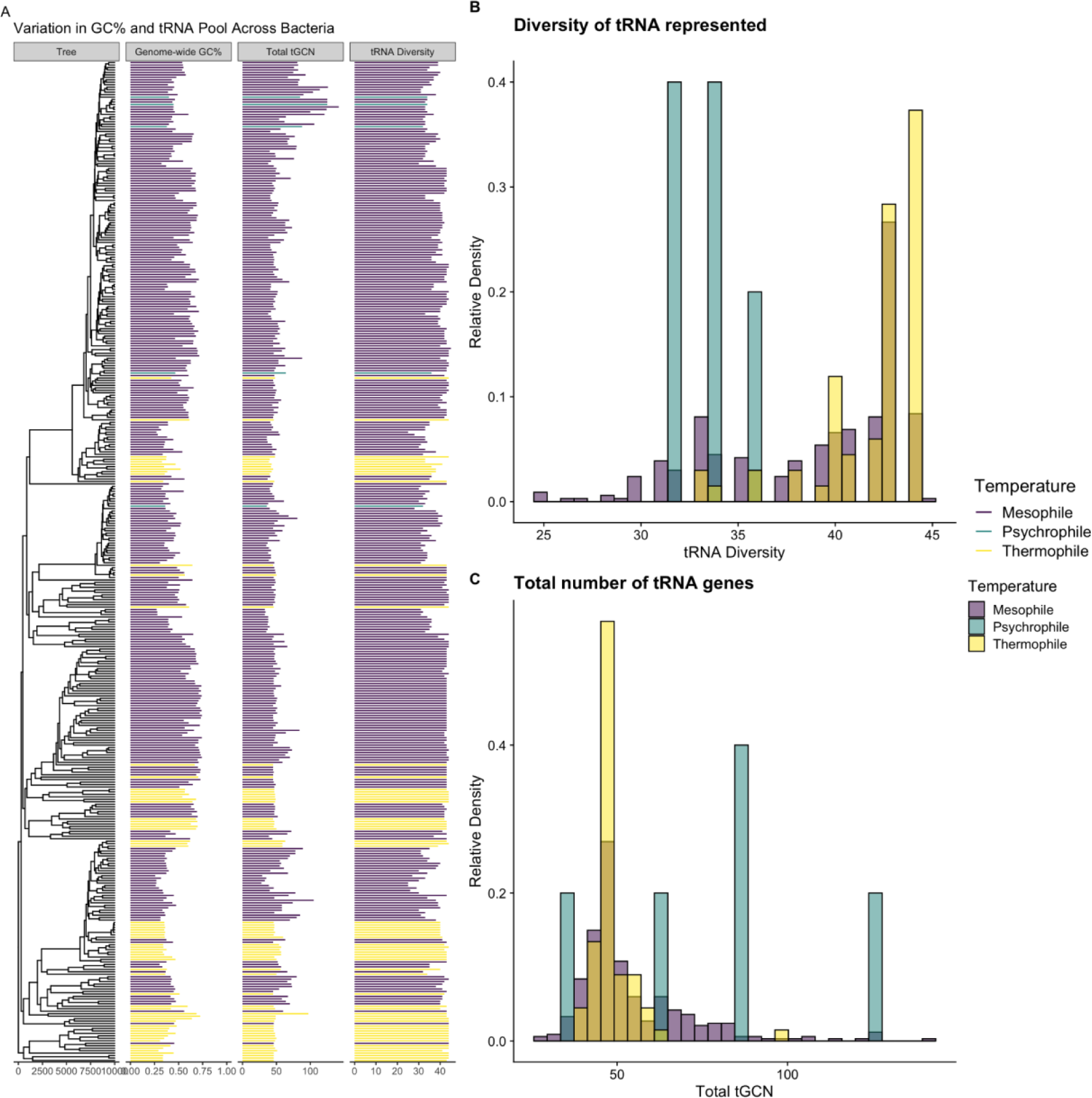
(A) Variation in genomic GC% and the tRNA pool vary across the species considered, as well as the evolutionary relationships of the species. Not all organisms studied are represented in the tree. (B) Distribution of total number of tRNA genes across all species for which data was available. (C) Distribution of the diversity of anticodons represented in tRNA genes across all species for which data was available.

We performed phylogenetic regressions to investigate the relationship between genomic GC%, tRNA diversity, total number of tRNA genes (total tGCN), and missense error rates *ϵ*_*M*_ across bacteria. These analyses indicate that higher GC% is associated with higher tRNA diversity, and both higher GC% and tRNA diversity are associated with higher missense error rates as determined by the median *ϵ*_*M*_ for each species (Figure 2). Interestingly, we observe that the relationship between GC% and tRNA diversity appears to be primarily driven by tRNA that recognizes GC-ending codons (not considering possible wobble) (Figure 3A). The representation of tRNA with anticodons recognizing codons ending in A or T/U varies with with GC%, but only weakly (Figure 3B). Notably, the presence/absence of tRNA with anticodons of the form CNN and GNN (where N is any of the 4 nucleotides) was previously observed to be variable across bacteria (Wald & Margalit 2014). Additionally, tRNA with anticodons of the form ANN are largely avoided due to modification to inosine (I), which could match with codons NNC, NNA, or NNU, increasing the potential for missense errors (Wald & Margalit 2014). As an NNC codon can only be decoded by an anticodon of the form GNN or INN (the latter of which may be largely avoided), and an NNG codon can only be decoded by an anticodon of the form CNN or UNN (the latter of which tRNA modifications can result in recognition of all possible codons), it is unsurprising that the diversity of tRNA recognizing GC ending codons scales with GC content.. A summary of model comparisons can be found in the Supplemental Material (Table S1).

**Figure 2.**
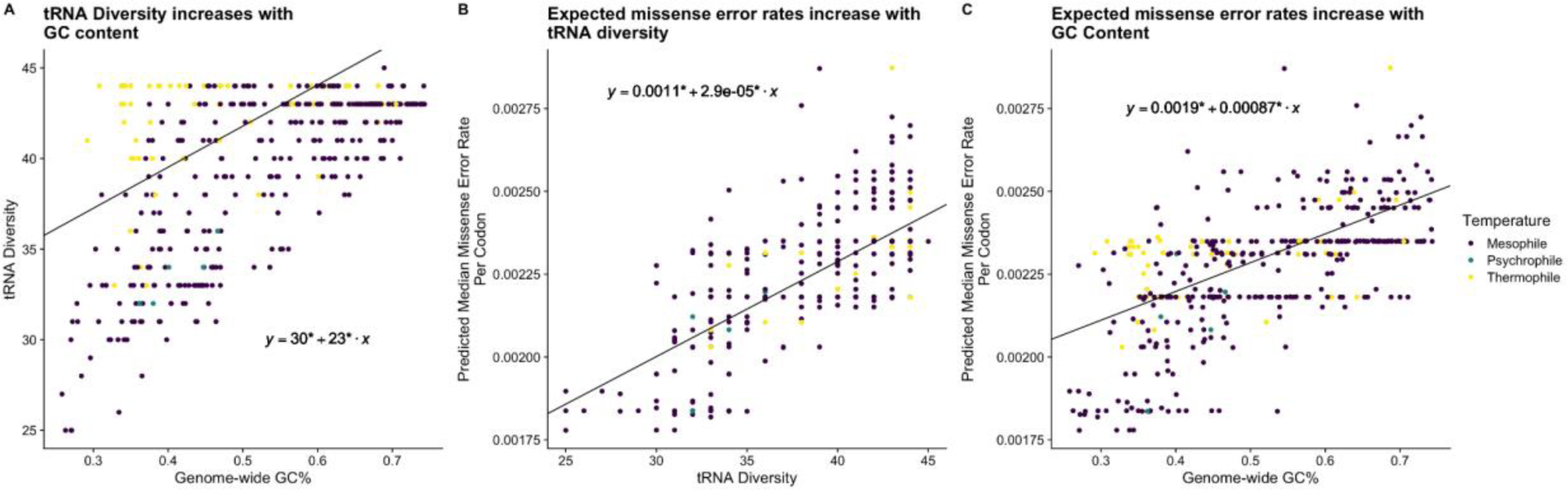
Phylogenetic regressions across all bacteria represented by solid black lines. * indicate parameter estimates significantly (p < 0.05) different from 0. (A) Demonstration of a positive relationship between tRNA diversity and GC% across bacteria (B) Demonstration of a positive relationship between tRNA diversity and predicted median *ϵ*_*M*_(C) Demonstration of a positive relationship between the median missense error rate for each species and GC% across bacteria

**Figure 3.**
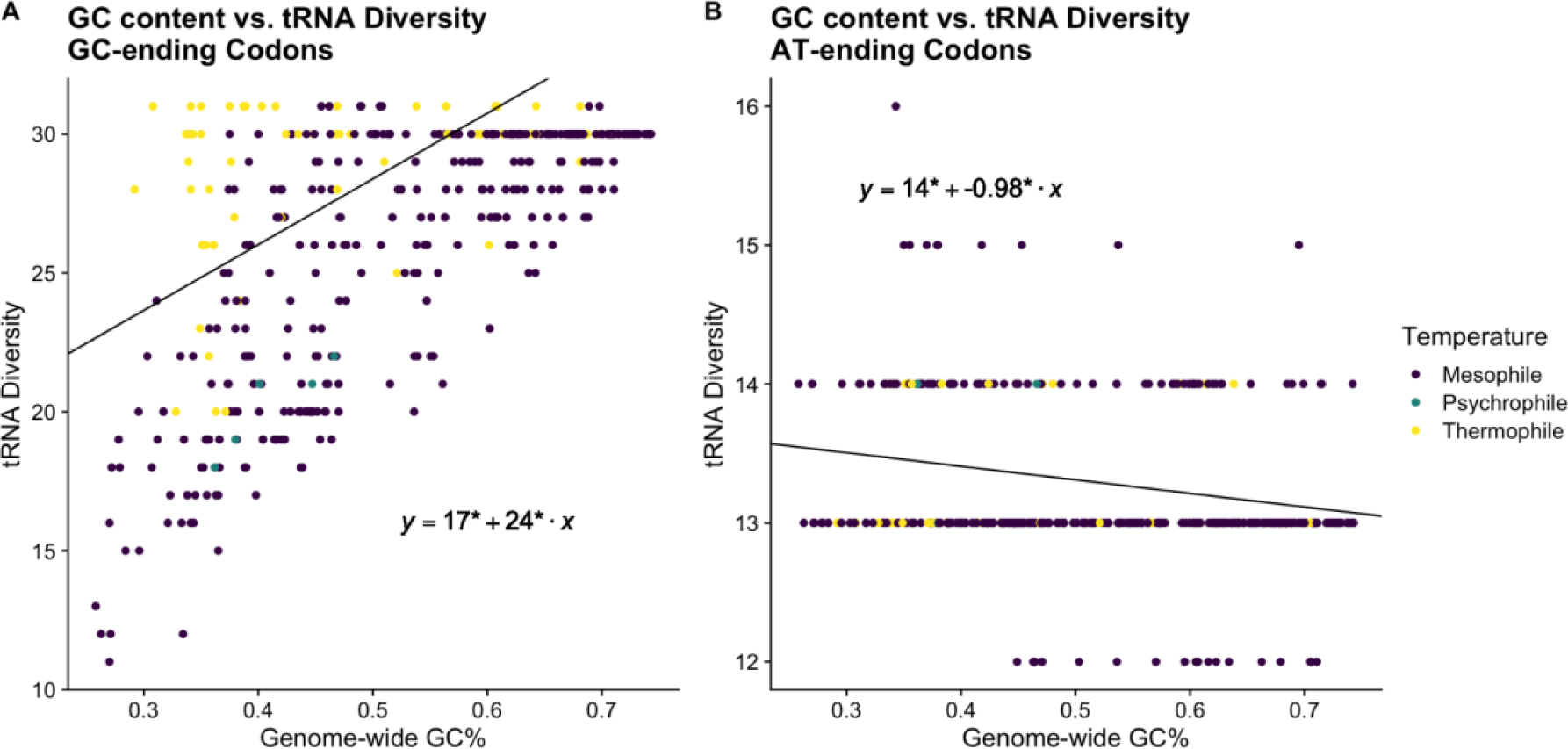
Differences in the evolution of tRNA anticodons recognizing GC vs. AT ending codons. Phylogenetic regressions performed across all bacteria. * indicate statistical significance (p < 0.05) of parameter estimates. (A) Relationship between genome-wide GC% and the diversity of tRNA recognizing GC codons. (B) Same as in (A), but for tRNA recognizing AT-ending codons.

### Testing for systematic differences in the tRNA pool related to environment

Previous work found that thermophiles have a greater tRNA diversity compared to mesophiles and psychrophiles (Satapathy et al. 2010). This analysis was based on a limited set of bacteria and failed to control for the shared ancestry of the species. Controlling for the effects of GC content in our phylogenetic regression, we observe that tRNA diversity is, on average, higher in thermophiles. Thermophiles were found to have approximately 1 to 2 more anticodons represented in the tRNA pool, on average, compared to mesophiles with a similar GC% (Table 1. *β*_*Thermo*_ = 1.77, *p* < 0.0001). We note that the use of a linear model to fit the bounded discrete tRNA diversity values may cause problems in our interpretation. In contrasts, we find no evidence that thermophiles have fewer total tRNA genes, on average, compared to mesophiles with a similar GC%. (Table S2. *β*_*Thermo*_ = −2.87, *p* = 0.15). Although the slope estimate for psychrophiles was positive (Table S2. *β*_*Psychro*_ = 3.70, *p* = 0.36), it was not statistically significant, such that we have insufficient evidence to claim that psychrophiles have a greater number of tRNA genes than their mesophilic counterparts, contrary to previous claims. However, this is likely due to a lack of statistical power, as we only had 5 psychrophiles in our final set of bacteria. Unlike with tRNA diversity, we found no evidence that GC content had a significant relationship with the total number of tRNA found in a genome (Table S2. *β*_*GC*_ = *3.9649*, *p* = 0.56).

**Table 1.**
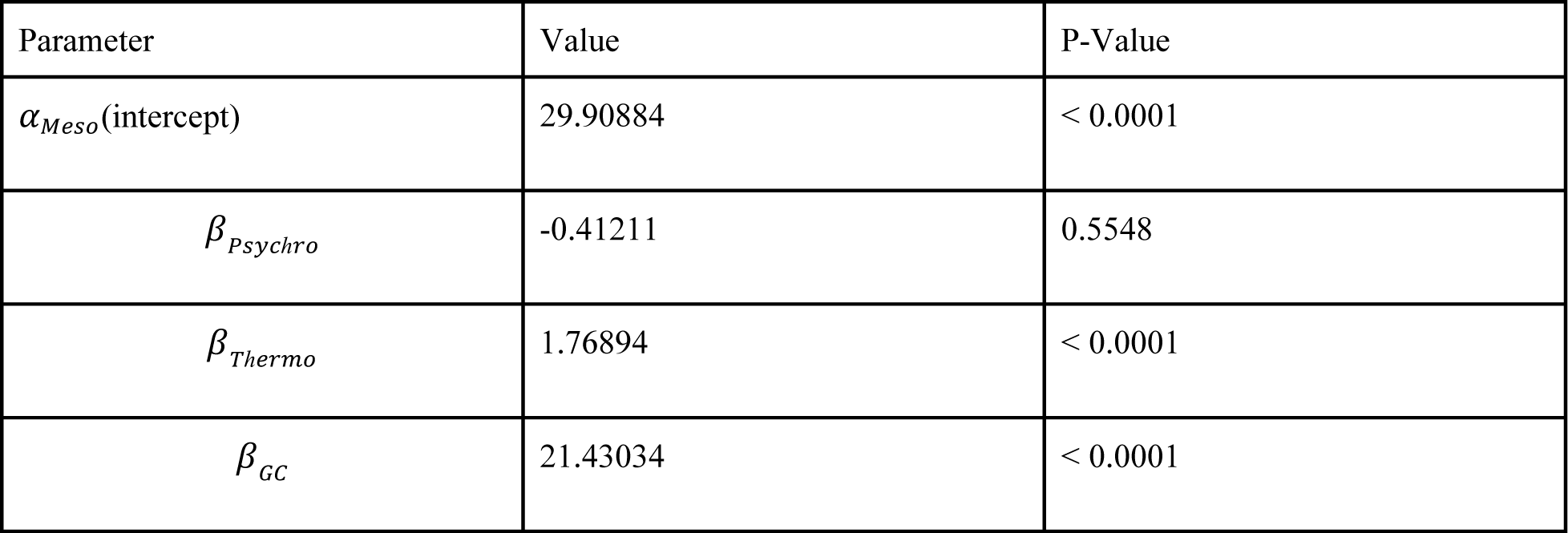
Phylogenetic regression parameter estimates when comparing tGCN diversity across mesophiles, thermophiles, and psychrophiles while taking GC% into account. This can be represented by the formula 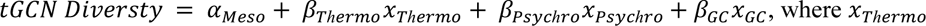 indicate if the bacteria is a thermophile or psychrophile (i.e., *x*_*Thermo/Psychro*_ = *1*, and 0 otherwise).. This means the slope estimates *β*_*Thermo*_ and *β*_*Psychro*_represent the mean value of tGCN diversity relative to a mesophilic bacteria. This regression assumed an OU model of trait evolution, which was 2.92 AIC units better than the same regression using a BM model of trait evolution.

| Parameter | Value | P-Value |
| --- | --- | --- |
| $\alpha_{Meso}$ (intercept) | 29.90884 | < 0.0001 |
| $\beta_{Psychro}$ | -0.41211 | 0.5548 |
| $\beta_{Thermo}$ | 1.76894 | < 0.0001 |
| $\beta_{GC}$ | 21.43034 | < 0.0001 |

Based on the general relationship between tRNA diversity and missense error rates observed across all bacteria, the increased diversity of thermophilic tRNA pools is expected to be associated with an increased missense error rate compared to psychrophiles and mesophiles. Our results support this hypothesis, as thermophiles were found to have a median missense error rate approximately *9* × *10*^−^*^5^*/s higher compared to mesophiles with the same GC% (Table S3. *β*_*Thermo*_ = *0*.*00009*, p < 0.0001). We emphasize missense error rate estimates are based on a theoretical model of translation largely based on measurements performed using *E. coli*. The safest interpretation of our missense error rate estimates is the error rates expected in *E. coli* given its tRNA pool looked like a particular species. Similar to our hypothesis that wobble may be less efficient in thermophiles due to the higher temperatures, missense errors may also be less likely at higher temperatures due to the instability in the codon-anticodon bonds. If increased temperatures do not offset the potential negative consequences of a more diverse tRNA pool, then the higher missense error rates may have implications for the evolution of protein synthesis in thermophiles.

First, we speculate that higher missense error rates in thermophiles could be compensated for via the evolution of more accurate ribosomes or changes to tRNA modifications. Previous work concluded that hyper-accurate ribosomes result in a substantial decrease in growth rates due to kinetic inefficiency (Ruusala et al. 1984). The impact of higher temperatures on reaction rates may help offset some of the negative consequences of more accurate ribosomes. Such arguments are similar to those attempting to explain the apparent weaker codon usage bias observed in thermophiles compared to mesophiles and psychrophiles: as higher temperatures increase reaction rates, the ability of natural selection to act on translation efficiency via codon usage is weakened (Vieira-Silva & Rocha 2010; Botzman & Margalit 2011). More in-depth studies of ribosome function in extremophiles are needed to empirically determine if differences in the elongation process systematically covary with environmental temperature.

Second, the higher missense error rates may have significant consequences for thermophilic proteome evolution. Thermophilic proteomes are known to contrast with their mesophilic counterparts. Thermophilic proteins generally accumulate amino acid substitutions at a slower rate than mesophilic proteins, hypothesized to be due to amino acid mutations being generally more destabilizing at higher temperatures (Zeldovich et al. 1 2007; Venev & Zeldovich 1 2018; Faure & Koonin 2015; Cherry 2010). As missense errors alter a protein sequence, missense errors could have a higher fitness burden in thermophiles. Thus, a higher missense error rate is expected to impose a greater fitness burden on thermophiles. A prominent hypothesis proposes natural selection against missense errors increases the frequency of “optimal” codons in both highly-expressed genes and at functionally important positions in the protein (Drummond & Wilke 2008; Yang et al. 1 2010). Although thermophiles generally have weakly biased codon usage, little work has examined codon usage at functionally important sites. Future work will focus on detecting signals of adaptive codon usage in thermophiles related to selection against missense errors.

## Conclusions

Here, we present evidence that extreme temperature environments help shape the evolution of the tRNA pool, a crucial feature of genome evolution and protein synthesis dynamics. As the tRNA pool is intimately tied with codon usage, deeper insight into the evolution of the tRNA pool will provide insight into how and why codon usage varies across species, particularly as it relates to the environment. Future research will also benefit from considering the presence and absence of tRNA modification enzymes, which can significantly impact protein synthesis (Manickam et al. 2016; Diwan & Agashe 2018; Nedialkova & Leidel 6 2015). We again emphasize that many of our analyses and hypotheses were extrapolated from observations in mesophiles. Comparative analyses of genome and proteome evolution as it relates to environmental temperature will greatly benefit from empirical research into the biochemistry and molecular biology of extremophilic organisms.

## Data availability

Scripts for recreating our analyses can be found at https://github.com/jainv0127/Shah_Labs

## Funding

This work was supported by the National Institute of Health-funded Rutgers INSPIRE IRACDA Postdoctoral Program [#GM093854, A.L.C].

## Acknowledgments

We would like to acknowledge Premal Shah for his helpful comments in the writing of this manuscript.

## Notes

https://github.com/jainv0127/Shah_Labs

### Competing Interest Statement

The authors have declared no competing interest.

### Summary of Updates

After submitting the original version, we discovered some bugs in our code. These (fortunately) had little impact on our biological conclusions. We have also updated the figures to be more color-blind-friendly.

