## supplemental for "Determining the effects of temperature on the evolution of bacterial tRNA pools"

### Supplemental Tables

| Phylogenetic Regression | Model | AIC | ΔAIC (relative to best) |
|---|---|---|---|
| GC vs. tRNA Diversity | BM | 1796.698 | 0 |
| | OU | 1797.452 | 0.754 |
| tRNA Diversity vs. missense error rate | BM | -5956.54 | 140.5 |
| | OU | -6097.04 | 0 |
| GC vs. missense error rate | BM | -5948.735 | 118.309 |
| | OU | -6067.044 | 0 |
| GC vs. tRNA Diversity (GC-ending codons) | BM | 1791.535 | 0 |
| | OU | 1793.672 | 2.137 |
| GC vs. tRNA Diversity (AT-ending codons) | BM | 653.397 | 77.831 |
| | OU | 575.566 | 0 |

Table S1. Model fit comparison of phylogenetic regressions based on either the Brownian Motion (BM) or Ornstein-Uhlenbeck (OU) models of trait evolution. Model comparisons are based on the Akaike Information Criterion (AIC). Models with the lower AIC are considered the better model.

| Parameter | Value | P-Value |
|---|---|---|
| $\alpha_{Meso}$(intercept) | 49.0393 | < 0.0001 |
| $\beta_{Psychro}$ | 3.9119 | 0.3373 |
| $\beta_{Thermo}$ | -2.9006 | 0.15 |
| $\beta_{GC}$ | 3.9649 | 0.5604 |

Table S2. Phylogenetic regression parameter estimates when comparing total tGCN across mesophiles, thermophiles, and psychrophiles while taking GC% into account. This can be represented by the formula $Total\ tGCN\ = \alpha_{Meso} + \beta_{Thermo}x_{Thermo} + \beta_{Psychro}x_{Psychro} + \beta_{GC}x_{GC}$, where $x_{Thermo}$ and $x_{Psycrho}$ indicate if the bacteria is a thermophile or psychrophile (i.e., $x_{Thermo/Psychro} = 1$, and 0 otherwise). This means the slope estimates $\beta_{Thermo}$ and $\beta_{Psychro}$represent the mean value of the total tGCN relative to a mesophilic bacteria. This regression assumed an OU model of trait evolution, which was 53 AIC units better than the same regression based on a BM model.

| Parameter | Value | P-Value |
|---|---|---|
| $\alpha_{Meso}$(intercept) | 0.0018 | < 0.0001 |
| $\beta_{Psychro}$ | -0.0000275 | 0.63090 |
| $\beta_{Thermo}$ | 0.0000547 | 0.01164 |
| $\beta_{GC}$ | 0.00089 | < 0.0001 |

Table S3. Phylogenetic regression parameter estimates when comparing the median missense error rates across mesophiles, thermophiles, and psychrophiles while taking GC% into account. We note this This can be represented by the formula $Median\ Missense\ Error\ Rate = \alpha_{Meso} + \beta_{Thermo} x_{Thermo} + \beta_{Psychro} x_{Psychro} + \beta_{GC} x_{GC}$, where $x_{Thermo}$ and $x_{Psycrho}$ indicate if the bacteria is a thermophile or psychrophile (i.e., $x_{Thermo/Psychro} = 1$, and 0 otherwise). This means the slope estimates $\beta_{Thermo}$ and $\beta_{Psychro}$represent the mean value of the median missense error rates relative to a mesophilic bacteria. This regression assumed an OU model of trait evolution, which was 124 AIC units better than the same regression based on a BM model.
